# Exact breakpoints of *In(1)w*^*m4*^ rearrangement

**DOI:** 10.1101/2021.12.08.471849

**Authors:** A. A. Solodovnikov, S. A. Lavrov

## Abstract

*In(1)w*^*m4*^ was known for decades as a classic example of position effect variegation-causing rearrangement and was mentioned in hundreds of publications. Nevertheless, the euchromatin breakpoint position of the rearrangement was not precisely localized. We performed nanopore sequencing of DNA from *In(1)w*^*m4*^ homozygous flies and determined the exact position of euchromatic (chrX:2767875) and heterochromatic breakpoints of the rearrangement.

## Introduction

*In(1)w*^*m4*^ inversion represents the first described case of heterochromatin position effect (Muller 1930). An inversion with breakpoints near the *white* gene and in the heterochromatin of the left arm of chromosome X (Figure 1, A) results in a mosaic inactivation of the *white* gene visible in the eyes of adult flies. The rearrangement is a classic model of PEV and has been mentioned in more than a hundred publications. In particular, in (Vogel *et al.*, 2009), the chromatin changes in the case of *In(1)w*^*m4*^-induces PEV were tracked using high-throughput methods (microarray hybridization and sequencing). In this paper, the euchromatin breakpoint position of inversion was determined with 1.5 kb accuracy (chrX:2766979-2768245, R6.22).

**Figure 1.**
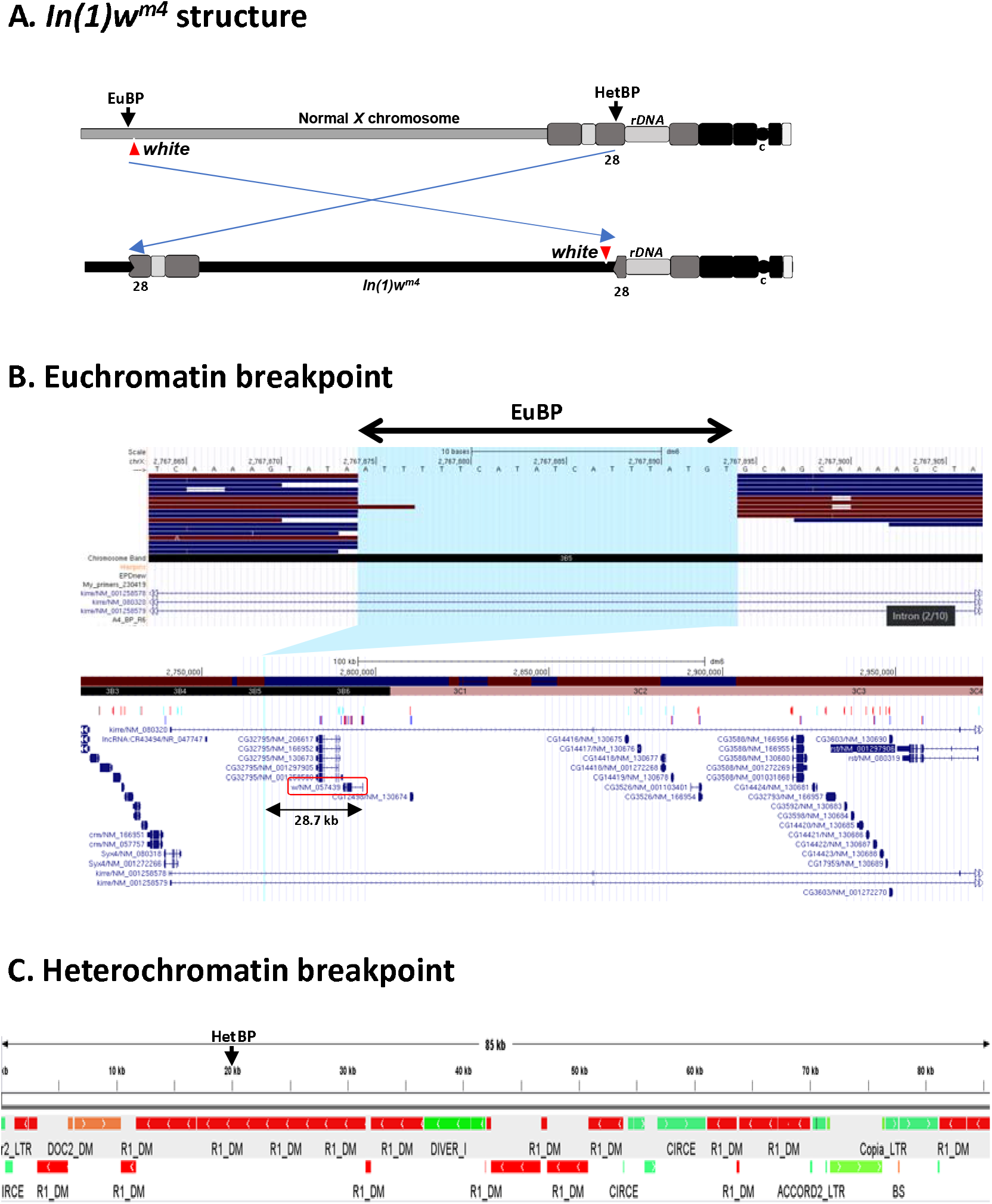
A. *In(1)w*^*m4*^ inversion scheme. EuBP – euchromatin breakpoint, HetBP – heterochromatin breakpoint, c – centromere, positions of h28 heterochromatin block and rDNA cluster are marked. Triangles denote the position of the *white* gene. B. Mapping of long reads to R6.22 *Drosophila* genome. The image is a collage of UCSC genome browser screenshots, the upper part is a close-up view of euchromatin breakpoint and reads mapped to this place, the lower part is an overview of 200 kb genomic region containing the breakpoint. The *white* gene is outlined, the arrow shows the distance between *white* TSS and the breakpoint. C. Heterochromatin breakpoint of *In(1)w*^*m4*^ inversion. The breakpoint in heterochromatin (HetBP) is in the element *R1* in the region, saturated with different types of mobile elements.

The heterochromatin breakpoint of *In(1)w*^*m4*^ was mapped to h28 block of X chromosome heterochromatin distal to rDNA cluster, and the sequence immediately near the breakpoint corresponds to R1 transposon (Appels *et al.*, 1982; Tartof *et al.*, 1984).

We decided to precisely map the positions of *In(1)w*^*m4*^ breakpoints to study the changes in chromatin organization immediately at the eu-heterochromatin border. The nanopore sequencing was applied to look deep into the heterochromatin beyond the breakpoint position. Nanopore sequencing is characterized by a significantly higher number of errors compared to traditional NGS approaches but has almost unlimited read length and, thus, extremely helpful in assembly of repeat-rich regions of heterochromatin.

## Materials and Methods

### DNA isolation

DNA from *In(1)w*^*m4*^ flies was extracted using proteinase K treatment – phenol-chloroform extraction – ethanol precipitation protocol. 100 flies were disrupted in 1 ml of TET buffer (100 mM Tris pH 8.5, 50 mM EDTA, 0.2% Triton X-100) in ice by Potter homogenizer. Sodium sarcosinate was added to 1.5% and proteinase K to 100 μg/μl, the suspension was gently mixed and incubated for 1 h at 50 °C. The mixture was extracted twice with phenol pH 8.0, once with the phenol-chloroform mix, and once with chloroform on Hula Mixer. The water phase after extraction was collected and sodium chloride was added to 100 mM. Then, 2 volumes of 96% EtOH were added and the tube slowly rotated until HMW DNA forms a tangle. DNA was washed once with 500 μl of 70% EtOH, dried, and dissolved in 100 μl of MilliQ water. DNA concentration was measured using Qubit fluorimeter (DS DNA broad range kit). Liquid handling at every stage was performed gently and using cut off pipette tips.

### MinIon sequencing

The sequencing library was prepared from 1.5 μg of DNA from homozygous *In(1)w*^*m4*^ flies according to the protocol recommended for Ligation Sequencing Kit (SQK-LSK109) from ONT (https://store.nanoporetech.com/us/media/wysiwyg/pdfs/SQK-LSK109/Genomic_DNA_by_Ligation_SQK-LSK109_-minion.pdf). The prepared library was loaded to MinIon R 9.4.1. flowcell and sequenced without basecalling until ~4 Gb of data was obtained.

### Data treatment

A set of .fast5 files from the sequencing run was basecalled on a standalone GPU-enabled server with Guppy Version 3.5.2 using dna_r9.4.1_450bps_hac profile. The resulting FASTQ files were loaded to the local Galaxy server for further processing. Quality checks using Nanostat (https://github.com/wdecoster/nanostat) and Nanoplot (https://github.com/wdecoster/NanoPlot) tools in Galaxy showed that ~2.4 Gb of reads (~20x genome coverage) with N50=14010 and Q>7 was received. Adapters were trimmed by Porechop (https://github.com/rrwick/Porechop) using default settings (reads with middle adapter were splitted).

Processed reads were mapped to r6.22 release of *D. melanogaster* genome using *Minimap2* software with Oxford Nanopore read to reference mapping profile (minimap2 -x map-ont) (Li 2018). The resulting .bam file was visualized in the local *UCSC Genome Browser* (Figure 1, B). FASTQ files with reads were also converted to FASTA using FASTQ to FASTA converter in Galaxy (Blankenberg *et al.*, 2010).

The euchromatin breakpoint was identified as a gap in aligned reads in the 2 kb region where *In(1)w*^*m4*^ breakpoint was located previously (Vogel *et al.*, 2009). The regions of 400 bp in size immediately upstream and downstream to the breakpoint (chrX:2767482-2767880 and chrX: 2767881-2768281) were blasted against sequencing reads in multifasta format to identify and extract reads overlapping breakpoint. 32 reads containing sequence near the presumptive breakpoint and unknown part corresponding to fused heterochromatin were identified. Two reads with the longest heterochromatin parts to the left and the right direction from the breakpoint were selected and their heterochromatin parts combined into one sequence. This sequence contains a region of heterochromatin ~80 kb in size, encompassing *In(1)w*^*m4*^ breakpoint (Figure 1, C).

The structure of heterochromatin near the *In(1)w*^*m4*^ breakpoint was analyzed using web application RepeatMasker (http://www.repeatmasker.org/cgi-bin/WEBRepeatMasker), which allows mapping of most types of *Drosophila* repeats. The output of RepeatMasker was manually converted to a .bed file, color information for different types of repeats added and the result visualized in IGV (https://software.broadinstitute.org/software/igv/) (Figure 1, C).

### Data availability

The sequences encompassing *In(1)wm4* breakpoints and 85 kb uninterrupted heterochromatin fragment are available in Supplementary. MinIon dataset with *In(1)w*^*m4*^ reads is available from SRA (SRR17055722).

## Results

### The results of sequences analysis are presented in Figure 1

There is a gap in a reads coverage in the genome region chrX:2766979-2768245, which, as it was shown previously (Vogel *et al.*, 2009), contains *In(1)w*^*m4*^ breakpoint. The position of the gap is chrX:2767875-2767894 according to R6.22 release of *D. melanogaster* genome (Figure 1, B). We used BLASTN to extract reads containing sequences immediately near this gap and investigate their composition. All these sequences contain euchromatin part fused to *Drosophila melanogaster* type I transposable element *R1Dm*. The breakpoint position in *R1* is 2200 according to GenBank X51968.1 *R1* sequence.

Eventually, the position of euchromatin breakpoint of *In(1)w*^*m4*^ is chrX:2767875-2767894, the inversion is accompanied by a deletion of 20 nucleotides of euchromatin, and the breakpoint is 28.7 kb distal to *white* TSS (Figure 1, B).

To identify sequences around the *In(1)w*^*m4*^ breakpoint in the heterochromatin, reads of maximum length containing the euchromatin regions closest to the breakpoint were extracted from the library of reads. Thus, the reads overlapping the breakpoint and containing ~80 kb of heterochromatin were detected. The structure of the heterochromatin was analyzed using the RepeatMasker. It was found that the heterochromatin breakpoint is in a region composed of fragments of different types of mobile elements (LINE (mainly R1) and LTR-containing), of different orientation and length. This type of organization resembles some piRNA-producing loci. Interestingly, the insertion of *I*-element in the h28 heterochromatin region leads to decreased reactivity in I-R hybrid dysgenesis system, pointing to the suppression of *I*-element transpositions (Dimitri *et al.*, 2005).

## Supporting information

Breakpoint flancs

Heterochromatic breakpoint elements

Heterochromatic breakpoint

## Acknowledgments

Authors are grateful to V. A. Gvozdev for his valuable comments and help in the preparation of this manuscript. The work was supported by the Russian Science Foundation (project No. 19-74-20178) and by the Russian Foundation for Basic Research (17-00-00282 KOMFI).

## Compliance with ethical standards

This article does not contain any studies involving animals or human participants performed by any of the authors.

